# Differences in Genetic Liability for Insomnia and Hypersomnia in Bipolar Disorder Subtypes

**DOI:** 10.1101/569376

**Authors:** Katie J S Lewis, Alexander Richards, Ganna Leonenko, Samuel E Jones, Hannah Jones, Katherine Gordon-Smith, Liz Forty, Valentina Escott-Price, Michael J Owen, Michael N Weedon, Lisa Jones, Nick Craddock, Ian Jones, Michael C O’Donovan, Arianna Di Florio

## Abstract

**Background:** Insomnia and hypersomnia are common in bipolar disorder (BD) but it is unclear whether genetic factors influence this association. Stratifying by bipolar subtypes could elucidate the nature of this association and inform research on sleep and BD. We therefore aimed to determine whether polygenic risk scores (PRS) for insomnia, daytime sleepiness and sleep duration differ according to bipolar subtypes (bipolar I disorder, BD-I or bipolar II disorder, BD-II).

**Methods:** In this case-control study, we used multinomial regression to determine whether PRS for insomnia, daytime sleepiness, and sleep duration were associated with risk of BD-I or BD-II when compared to each other and to controls. Cases (n=4672) were recruited within the United Kingdom from the Bipolar Disorder Research Network. Controls (n=5714) were recruited from the 1958 British Birth Cohort and UK Blood Service. All participants were of European ancestry. Clinical subtypes of BD were determined by semi-structured psychiatric interview (the Schedules for Clinical Assessment in Neuropsychiatry) and case notes.

**Results:** Within cases, 3404 and 1268 met DSM-IV criteria for BD-I and BD-II, respectively. Insomnia PRS was associated with increased risk of BD-II (RR = 1.14, 95% CI = 1.07-1.21, *P* = 8.26 × 10^−5^) but not BD-I (RR = 0.98, 95% CI = 0.94-1.03, *P* = .409) relative to controls. In contrast, sleep duration PRS was associated with increased relative risk of BD-I (RR = 1.10, 95% CI = 1.06-1.15, *P* = 1.13 × 10^−5^), but not BD-II (RR = 0.99, 95% CI = 0.93-1.06, *P* = .818). Daytime sleepiness PRS was associated with an increased risk of BD-I (RR = 1.07, 95% CI = 1.02-1.11, *P* = .005) and BD-II (RR = 1.14, 95% CI = 1.07-1.22, *P* = 3.22 × 10^−5^) compared to controls, but did not distinguish subtypes from each other.

**Conclusions:** Bipolar subtypes differ in genetic liability to insomnia and sleep duration. Our findings provide further evidence that the distinction between BD-I and BD-II has biological and genetic validity. This distinction will be critical in the selection of participants for future research on the role of sleep disturbance in BD.

## Introduction

Bipolar disorder (BD) is a severe mood disorder which impacts quality of life and wider society.^1, 2^ Insomnia (difficulty initiating and maintaining sleep) and hypersomnia (excessive daytime sleepiness or prolonged sleep duration) form part of the diagnostic criteria for depressive episodes.^3^ These sleep disturbances may persist in the inter-episode period,^4–9^ and are associated with significant distress and impairment.^4^ Furthermore, reduced sleep duration, a symptom of manic episodes, has been implicated as a prodrome and trigger of mania.^10–14^ These findings have initiated interventions that aim to improve wellbeing and reduce episode recurrence in BD by improving sleep.^15, 16^ An example is cognitive behavioural therapy for insomnia adapted for individuals with BD, with several randomised controlled trials piloted or currently in progress.^16–18^ Understanding the relationship between sleep and mood in BD is therefore important and could inform clinical interventions.

Recent genome-wide association studies (GWAS) of sleep traits such as insomnia, sleep duration and daytime sleepiness^19–24^ have offered researchers the opportunity to examine this relationship at the genomic level. Published studies have found no significant genetic correlations between bipolar disorder and insomnia using summary-level data^19, 21^ but these analyses may have been limited by a lack of individual bipolar phenotypic and genotypic data.

Bipolar disorder type 1 (BD-I) and type 2 (BD-II) are the primary clinical sub-types of BD,^3^ with evidence suggesting they differ in illness course, clinical features, and aetiology.^25–27^ BD subtypes may also differ in rates of insomnia; Steinan et al. examined sleep profiles in BD and found that during depressive episodes, insomnia symptoms were more prevalent among individuals with BD-II than those with BD-I.^28^ However, the authors cautioned that it was not possible to determine whether insomnia symptoms also differed in the inter-episode period (i.e. whether differences in trait liability to insomnia exist). Therefore, our first aim was to determine whether genetic liability for insomnia indexes different BD subtypes. Given a lack of evidence on whether prevalence of insomnia symptoms differs between BD subtypes in the inter-episode period, we had no prior hypothesis about which subtype, if any, would be genetically correlated with insomnia.

Insomnia is not the only sleep disturbance present in individuals with BD. Hypersomnia (defined as prolonged sleep duration or excessive daytime sleepiness^29^) is another common sleep disturbance in BD.^30, 31^ Hypersomnia is more common in bipolar depression than unipolar depression,^29, 32^ shows high recurrence across episodes of bipolar depression^33^ and persists in the inter-episode period.^31, 34, 35^ In the study of Steinan et al., hypersomnia was more prevalent in BD-I (during depression and euthymia) than in BD-II.^28^ Steinan et al., and other researchers have thus called for further understanding of the role of hypersomnia in BD and its underlying biology.^30, 36^

The two constructs that comprise hypersomnia, prolonged sleep duration or excessive daytime sleepiness, are often combined but there is evidence that the two are uncorrelated^37, 38^ and may differ in their relationships with bipolar disorder; a recent study found that excessive daytime sleepiness, but not sleep duration, predicted relapse to mania/hypomania.^39^ Therefore, our second aim was to investigate the genetic relationships between subtypes of hypersomnia (longer sleep duration and excessive daytime sleepiness) and BD subtypes.

In order to test the relationships between sleep and bipolar phenotypes, we adopted the polygenic risk score (PRS) method to estimate the burden of risk alleles associated with three sleep related phenotypes (insomnia, sleep duration and excessive daytime sleepiness) in people with BD-I, BD-II and in controls.^40, 41^ In secondary analyses, we conducted a two-sample Mendelian randomization (MR) study to explore whether sleep phenotypes increase risk of BD-I or BD-II.

## Method

### Sample Recruitment

#### Bipolar Disorder

Participants with BD (n=4672) were recruited within the United Kingdom by the Bipolar Disorder Research Network (bdrn.org).^42^ All participants self-reported as white, were unrelated, aged 18 years or over and recruited systematically (e.g. community mental health teams) or non-systematically (e.g. via websites, radio adverts or voluntary groups such as Bipolar UK). Participants were excluded if they had affective illness experienced solely in response to alcohol or substance misuse, or affective illness secondary to medical illness or medication. Participants provided written informed consent. The study had ethical approval from the West Midlands Multi-Centre Research Ethics Committee in addition to local research and development approval by UK National Health Service Trusts and Health Boards.

#### Controls

Controls (n=5,714) were recruited via the UK Blood service and the 1958 Birth Cohort (58C). Characteristics and recruitment of these samples has been described previously.^43^ All controls self-reported as white.

### Measures

Bipolar disorder cases were assessed using the Schedules for Clinical Assessment in Neuropsychiatry interview.^44^ Information from this interview was combined with clinical case note data to make lifetime best-estimate DSM-IV diagnoses. Inter-rater reliability for DSM-IV diagnoses was good, with a mean kappa statistic of 0.85.

### Genotyping, Quality Control, and Imputation

Genotyping was conducted on Affymetrix GeneChip 500K Mapping Array Set, Illumina Omni Express Array, and Illumina PsychChip. Strict quality control (QC) was performed separately on batches from each platform before merging (Supplementary Note 1). QC was conducted using PLINK version 1.9 software^45^ in which single nucleotide polymorphisms (SNPs) were excluded if the minor allele frequency (MAF) was less than 0.01, if SNPs deviated from Hardy-Weinberg Equilibrium (HWE) at *P* ≤ 10^−6^ or call rate < 98%. Individuals were excluded from the sample if they had increased or decreased heterozygosity of |F| > 0.1, a discrepancy between their genotypic and reported sex, genotype call rate < 98%, high pairwise relatedness (pi-hat _>_ 0.2) or did not cluster with European population samples in principal component analysis of 2000 participants from 19 populations of the 1000 Genomes Project.^46^

After QC, data for each platform were phased using SHAPEIT version 3.4.0.1023^47^ and imputed using IMPUTE2^48^ with the 1000 Genomes Project reference panel (phase 3). Imputed data were converted to the most probable genotypes (probability ≥ 0.9) with additional SNPs excluded if the imputation INFO score was <0.8, MAF<0.01 or HWE *P* <1×10^−6^). Imputed data were merged on shared SNPs. SNPs which showed large differences in frequency between samples on different genotyping platforms were removed (see Supplementary Note 1).

Control genotypic data were processed using the same method for QC, phasing and imputation as outlined for cases. After merging and removing SNPs with strand ambiguity, the final dataset used for analyses comprised 4,672 cases and 5,714 controls with 1,900,924 imputed SNPs.

### Principal Component Analysis

PLINK version 1.9 software (Chang et al., 2015) was used to conduct principal component analysis on the clumped dataset. Eigenvectors for the first 10 principal components were included in all association analyses in order to control for potential confounding from population structure.

### Discovery Datasets for Insomnia, Sleep Duration and Daytime Sleepiness

The discovery datasets were GWAS summary statistics for insomnia,^20^ sleep duration^23^ and daytime sleepiness^24^ that were conducted in participants recruited to UK Biobank.^49^ Sleep phenotypes were assessed using touchscreen questions. To assess insomnia symptoms, participants were asked, “Do you have trouble falling asleep at night or do you wake up in the middle of the night?” with responses “never/rarely,” “sometimes,” “usually” and “prefer not to answer.” The insomnia GWAS was conducted in 236,163 participants who answered “usually” (cases) or “never/rarely” (controls). The sleep duration GWAS was conducted in 448,609 participants who were asked, “About how many hours sleep do you get in every 24 hours? (please include naps)”. Responses were in hour increments and were analysed as a continuous variable. The daytime sleepiness GWAS was conducted in 452,071 participants and assessed using the question, “How likely are you to doze off or fall asleep during the daytime when you don’t mean to? (e.g. when working, reading or driving),” with responses “never/rarely,” “sometimes,” “often,” or “all the time,” analysed on a 1-4 scale.

### Polygenic Risk Scores

We generated polygenic risk scores (PRS) using PLINK version 1.9^45^ in PRSice.^50^ Imputed genotypes were clumped for linkage disequilibrium (LD; window 500kb, r^2^= 0.2) and SNPs most significantly associated with sleep traits (i.e. insomnia, sleep duration and daytime sleepiness) were retained. Clumping resulted in the retention of 92085, 92096 and 91950 SNPs for daytime sleepiness, sleep duration and insomnia respectively. After clumping, PRS for sleep traits were generated using PRSice^50^ at p-value thresholds p ≤ 1, p ≤ 0.5, p ≤ 0.2, p ≤ 0.1, p ≤ 0.05, p ≤ 0.01, p ≤ 0.001 and converted to z-scores. This range of *P*-value thresholds was chosen in the absence of an independent sample from which to determine the most predictive *P*-value threshold.

### Statistical analysis

Data analysis was conducted in R version 3.33. We performed multinomial logistic regression analyses examining associations between PRS for insomnia, sleep duration, and daytime sleepiness (at the range of p-value thresholds described above) and bipolar subtypes (BD-I or BD-II) vs. controls. All analyses were adjusted for sex and 10 population principal components. In sensitivity analyses, we performed direct comparisons between the BD subtypes using logistic regressions that corrected for age, sex and 10 population principal components. Results are reported at the *P*-value thresholds that showed the most significant results with a False Discovery Rate correction applied.

### Mendelian Randomisation Analyses

Where we observed significant associations between sleep phenotype PRS and BD subtypes, we conducted follow-up two-sample MR studies to test whether sleep phenotypes (exposures) were causally related to BD subtypes (the outcome). MR is a causal inference method that uses genetic variants as instrumental variables for an exposure of interest. MR relies on three assumptions: (i) the genetic variants must be strongly associated with the exposure, (ii) genetic variants should not be associated with confounders of the exposure-outcome relationship, and (iii) the genetic variants should only be associated with the outcome through the exposure in question.^51^ We used genome-wide significant SNPs as genetic instruments for the sleep phenotypes. Instrument-exposure effects were taken from the sleep trait GWAS summary statistics and instrument-outcome effects were taken from BD-I and BD-II GWAS summary statistics.^52^ Four MR methods were used to assess a causal association between sleep phenotypes and bipolar subtypes: inverse variance weighted,^53^ weighted median,^54^ weighted mode^55^ and MR Egger^53^ regression methods. To test for evidence of pleiotropy, we examined the intercept of MR Egger Regressions,^53^ and the Cochrane Q and Rucker’s Q statistics.^56, 57^ Data pruning, harmonization and analyses were conducted in R version 3.33 using the ‘TwoSampleMR’ R package.^58^

## Results

### Sample Description

Within BD cases, 67% were female with a median age of 46 years (range 18-89 years). 3404 met criteria for BD-I and 1268 met criteria for BD-II. Within controls, 49% were female.

### Correlations between PRS for Sleep Traits

Across all PRS *P*-value thresholds, insomnia PRS was negatively associated with sleep duration PRS (*r* range = −0.17 to −0.30, *P* < 1 × 10^−4^) and positively associated with daytime sleepiness PRS (*r* range = 0.04 to 0.10, *P* range .0069 to < 1 × 10^−4^). Sleep duration PRS was negatively associated with daytime sleepiness PRS (*r* range = −0.03 to −0.00) but this association was not significant across all thresholds (*P* range .028 to .916).

### Associations between PRS for Sleep Traits and BD Subtypes

Results for multinomial regressions at the most significant PRS *P*-value thresholds (with corrected *P-*values) are summarised in Table 1. Results for multinomial and logistic regressions at other PRS *P*-value thresholds (*P*_T_) are provided in Supplementary Tables 2-7.

**Table 1.**
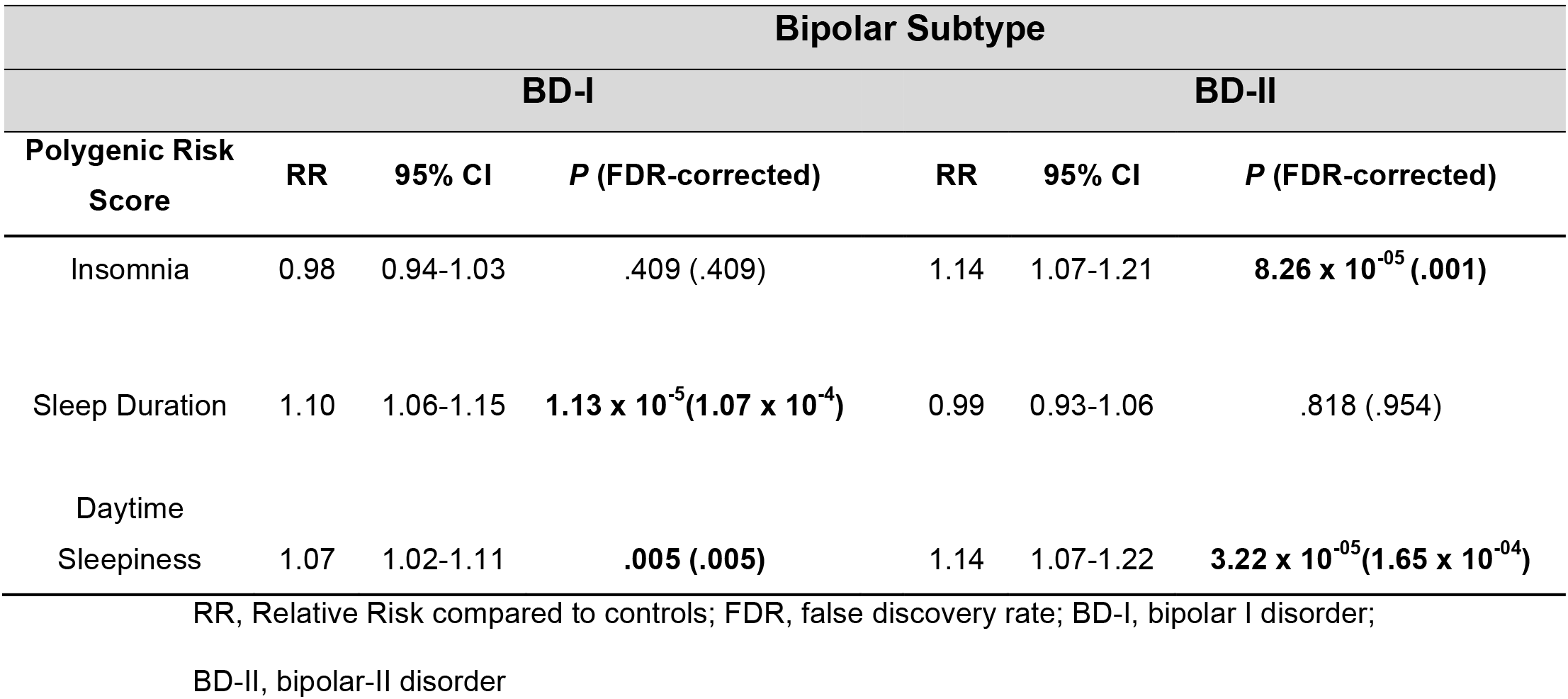
Summary of associations between polygenic risk scores (PRS) for sleep duration, insomnia and daytime sleepiness and bipolar subtypes compared to controls.

| Polygenic Risk Score | Bipolar Subtype |  |  |  |  |  |
| --- | --- | --- | --- | --- | --- | --- |
|  | BD-I |  |  | BD-II |  |  |
|  | RR | 95% CI | *P* (FDR-corrected) | RR | 95% CI | *P* (FDR-corrected) |
| Insomnia | 0.98 | 0.94-1.03 | .409 (.409) | 1.14 | 1.07-1.21 | **8.26 x 10<sup>-05</sup> (.001)** |
| Sleep Duration | 1.10 | 1.06-1.15 | **1.13 x 10<sup>-5</sup> (1.07 x 10<sup>-4</sup>)** | 0.99 | 0.93-1.06 | .818 (.954) |
| Daytime Sleepiness | 1.07 | 1.02-1.11 | **.005 (.005)** | 1.14 | 1.07-1.22 | **3.22 x 10<sup>-05</sup> (1.65 x 10<sup>-04</sup>)** |

RR, Relative Risk compared to controls; FDR, false discovery rate; BD-I, bipolar I disorder; BD-II, bipolar-II disorder

#### Associations between PRS for Insomnia and BD Subtypes

Multinomial regressions comparing BD subtypes to controls revealed that insomnia PRS was significantly associated with a decreased risk of BD-I at PRS *P*_T_ of *P* ≤ 1 and *P* ≤.5 but significant associations were not seen at other *P*-value thresholds (Supplementary Table 2). At all *P*-value thresholds, insomnia PRS was significantly associated with an increased relative risk of BD-II (RR = 1.14, 95% CI = 1.07-1.21, *P* = 8.26 × 10^−05^, *P*_FDR_ = .001 [PRS *P*_T_ ≤ .001]). Results at *P*_T_ ≤ .001 are shown in Figure 1. In direct tests, insomnia PRS significantly predicted BD-II compared to BD-I (OR = 1.14, 95% CI = 1.07-1.22, *P* = 6.81 × 10^−5^[PRS *P*_T_ ≤ .001] Supplementary Table 5).

**Figure 1.**
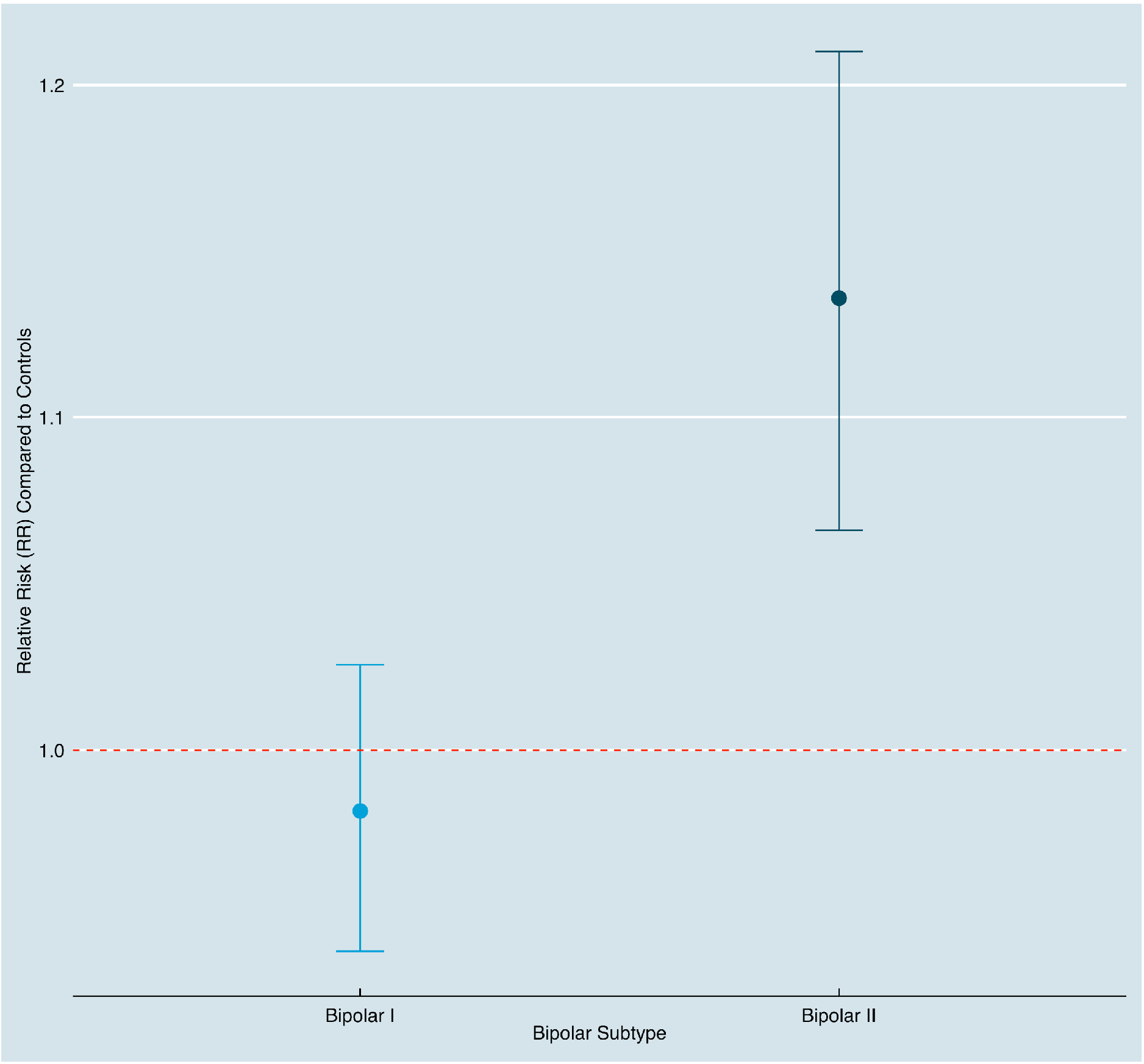
Relative risk (RR) of bipolar subtypes vs. controls predicted by polygenic risk for insomnia (PRS *P*-value threshold ≤ .001). Error bars indicate 95% confidence intervals.

#### Associations between PRS for Sleep Duration and BD Subtypes

At all PRS *P*-value thresholds, multinomial regression comparing BD subtypes to controls revealed that sleep duration PRS was associated with a significant increased relative risk of BD-I (RR = 1.10, 95% CI = 1.06-1.15, *P* = 1.13 × 10^−5^, *P*_FDR_ = 1.07 × 10^−4^ [PRS *P*_T_ ≤ 1] see Supplementary Table 3). Associations between sleep duration PRS and relative risk of BD-II were not significant at any PRS *P*-value threshold (Supplementary Table 2). Results at *P*-value threshold ≤ 1 are shown in Figure 2. Direct comparisons between BD-I and BD-II revealed that sleep duration PRS was significantly associated with BD-I (OR = 1.11, 95% CI = 1.04-1.19, *P* = .002, [PRS *P*_T_ ≤ 1] Supplementary Table 6).

**Figure 2.**
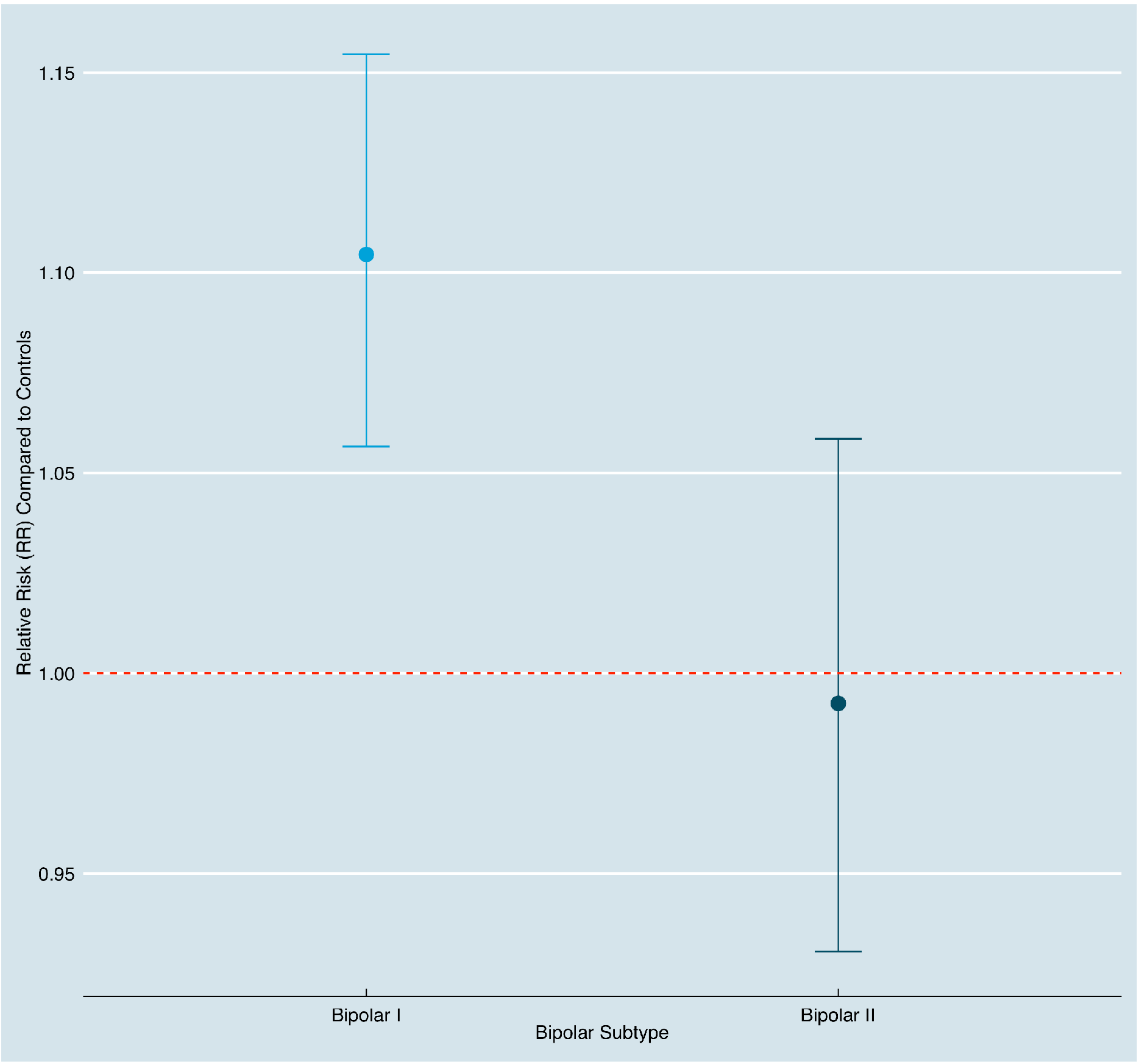
Relative risk (RR) of bipolar subtypes vs. controls predicted by polygenic risk for sleep duration (PRS *P*-value threshold ≤ 1). Error bars indicate 95% confidence intervals.

#### Associations between PRS for Daytime Sleepiness and BD Subtypes

Compared to controls, daytime sleepiness PRS was associated with an increased relative risk of BD-I and BD-II at all PRS *P*-value thresholds (except *P*_T_ < .001, see Supplementary Table 4). Results at PRS *P*_T_ ≤ .05 are shown in Table 1 and Figure 3. Direct comparisons between BD subtypes were not significant following correction for multiple testing (Supplementary Table 7).

**Figure 3.**
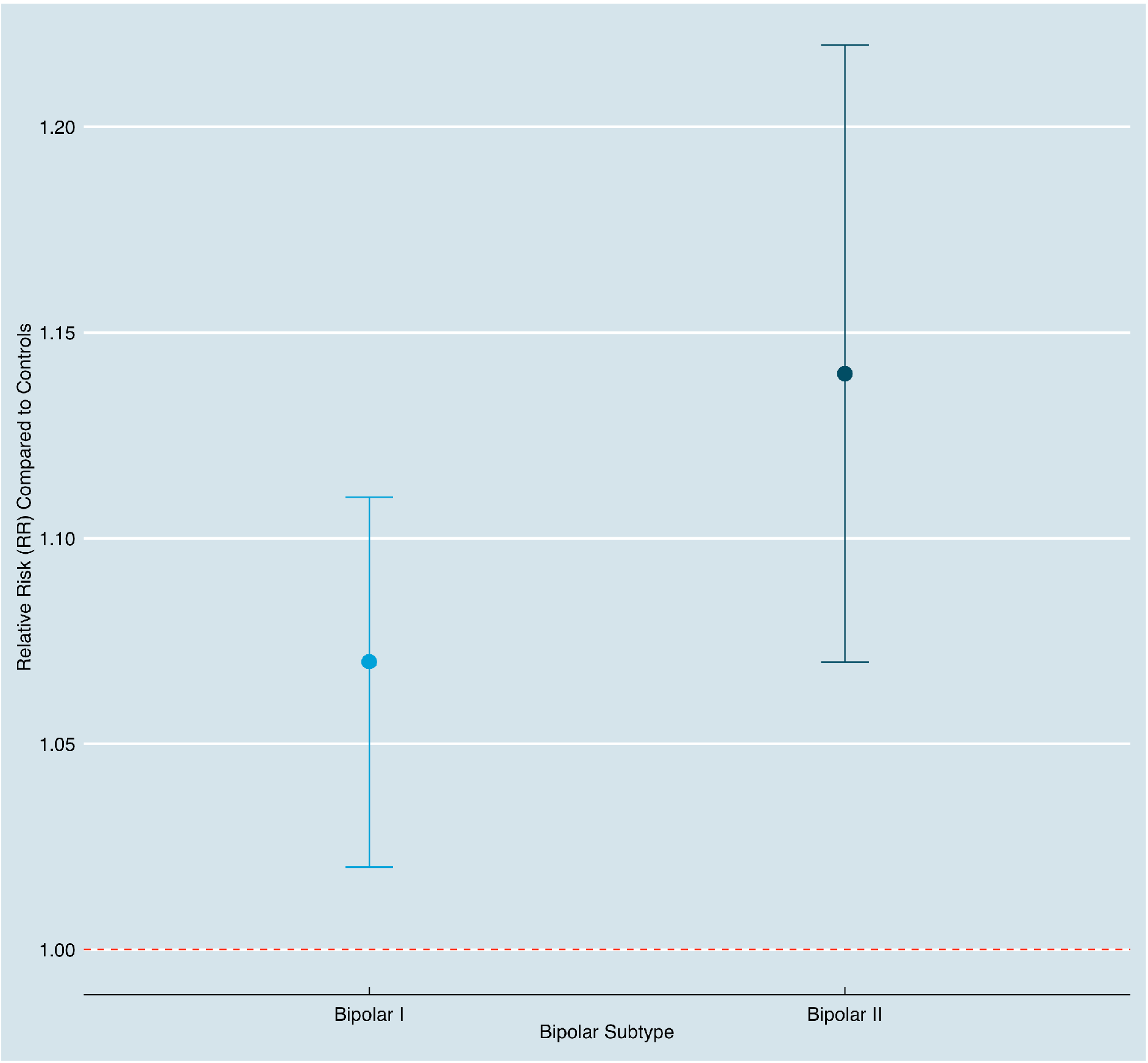
Relative risk (RR) of bipolar subtypes vs. controls predicted by polygenic risk for daytime sleepiness (PRS *P*-value threshold ≤ .05). Error bars indicate 95% confidence intervals.

### Mendelian Randomisation Analyses

Across all MR methods, we did not find strong evidence of a causal association between insomnia and BD-II or sleep duration and BD-I although the direction of effect was consistent when assessing a causal effect of insomnia on BD-II (Table 2). Although MR Egger intercepts were not significantly different from 0 for either analysis (*P* = 0.504 for insomnia to BD-II and *P* = 0.362 for sleep duration to BD-I), the Cochrane Q and Rucker Q statistics indicated significant heterogeneity in effect estimates (Table 2) possibly due to horizontal pleiotropy.

**Table 2.** Results of Two-Sample MR studies examining causal associations between insomnia and BD-II and sleep duration and BD-I.

| MR Method | Insomnia - BD-II (37 SNPs) |  |  | Sleep Duration - BD-I (56 SNPs) |  |  |
| --- | --- | --- | --- | --- | --- | --- |
|  | log(OR) | SE | *P* | log(OR) | SE | *P* |
| Inverse variance weighted | 0.256 | 0.149 | 0.087 | 0.396 | 0.209 | 0.059 |
| Weighted median | 0.355 | 0.183 | 0.052 | 0.245 | 0.219 | 0.264 |
| Weighted mode | 0.410 | 0.300 | 0.180 | 0.005 | 0.361 | 0.990 |
| MR Egger | 0.577 | 0.498 | 0.255 | -0.274 | 0.757 | 0.719 |
| Q Statistic | Q | df | *P* | Q | df | *P* |
| Rucker's Q | 53.33 | 35 | 0.0243 | 126.92 | 54 | **8.33 x 10<sup>-8</sup>** |
| Cochran Q | 54.03 | 36 | 0.0272 | 128.91 | 55 | **7.18 x 10<sup>-8</sup>** |

MR, Mendelian Randomisation; BD-I, bipolar I disorder; BD-II, bipolar-II disorder; df, degrees of freedom.

## Discussion

BD is heterogeneous in symptom presentation and most likely in the mechanisms that underlie these presentations. Genetics can help refine diagnostic groups that share a similar aetiology.^59^ Here, we provide the first evidence that genetic liability to insomnia and longer sleep duration differs according to bipolar subtypes.

Genetic liability to insomnia as indexed by PRS was associated with increased relative risk of BD-II compared to controls, and distinguished individuals with BD-II from those with BD-I. The stronger association between insomnia PRS and BD-II may explain non-significant genetic correlations between BD and insomnia in previous research,^19, 21^ as individuals with BD-II are usually underrepresented in BD GWAS (e.g. only 11% in a recent study^60^).

We used sleep duration and daytime sleepiness PRS as proxies for genetic liability to hypersomnia. Sleep duration PRS was associated with increased relative risk of BD-I but not BD-II, and was significantly more strongly associated with BD-I than BD-II when in a direct comparison. In contrast, daytime sleepiness PRS did not significantly distinguish bipolar subtypes. Daytime sleepiness PRS may be a proxy for insomnia, as daytime sleepiness can be induced by insomnia^61^ and we observed significant positive correlations between insomnia and daytime sleepiness PRS. Our results support existing research on the importance of hypersomnia in BD^30, 36^ (and in BD-I in particular^28^) and provide further evidence that hypersomnia is not a unitary construct^37, 38^

Genetic propensity to longer sleep duration could increase BD-I vulnerability by increasing the length of sleep needed. Longer sleep durations are generally in practice more difficult to obtain than short sleep durations, which could increase the likelihood of manic episodes being triggered by a sub-optimal number of hours slept. This could explain why individuals with BD-I are more likely than those with BD-II to report sleep loss triggering episodes of high mood.^12^

In contrast, increased genetic risk for insomnia in people with BD-II may reflect the known genetic associations between BD-II and depression.^52^ Both depression and insomnia are associated with negative attentional biases^62^ and are significantly genetically correlated.^19, 21, 63^ As such, GWAS of insomnia may be identifying genetic variants associated with being prone to negative affect.

The results of our MR analyses do not support causal associations between insomnia and BD-II or sleep duration and BD-I. However, we observed significant heterogeneity in our genetic instruments, thereby violating the third assumption of MR and potentially biasing our results. Therefore, whilst insomnia and sleep duration could be useful clinical stratifiers, there is currently insufficient evidence to conclude a causal role.

### Implications

Clinical and biological heterogeneity combined with a classification that is not grounded in biology are obstacles to advancing BD research. We provide evidence of heterogeneity in genetic propensity to sleep traits within BD, highlighting differences in the way that sleep-related genetic factors are associated with BD subtypes. This adds to previously published work on stratification in BD^64^ and work suggesting different aetiological factors influence the two conditions.^25–27^

Our results suggest that clinical trials of sleep interventions should stratify participants by clinical subtype and genetic liability to insomnia/hypersomnia. Future work should explore which factors drive the differences in genetic liability for insomnia/sleep duration between BD subtypes.

### Strengths and Limitations

This study was conducted on the world’s largest single cohort of BD with genotypic and phenotypic data. Phenotypic data were collected using face-to-face semi-structured interviews and case notes with high interrater reliability. We were therefore able to explore genetic associations using individual-level genetic data, which provided more granularity than summary statistics.

Our study has several limitations. First, potential recruitment bias in our BD sample may have reduced its representativeness and influenced our results.^65^ Second, we were unable to adjust for additional variables (e.g. age, education) because these were unavailable for controls. Second, our index of hypersomnia is imprecise, as the available GWAS summary statistics measured sleep duration as total hours slept;^23^ previous work suggests that hypersomnia is better characterised by total time in bed.^38, 66, 67^ Finally, variants associated with insomnia/hypersomnia at age 40-69 years (the age of the UK Biobank sample^49^) may differ from those associated in childhood or early adulthood. This may have increased our type 2 error rate as the majority of patients with BD experience the first onset of impairing symptoms in adolescence or early adulthood.^68^ Genetic risk for insomnia/hypersomnia that manifests during or prior to early adulthood may be more strongly associated with BD than those associated with mid-life insomnia. Our results should be replicated using future sleep trait GWAS in younger samples that are of sufficient size for PRS analysis.

## Conclusions

To our knowledge, this is the first study to explore whether genetic liability for sleep traits is associated with clinical strata of individuals with BD. Future work should explore the potential mechanisms underlying differences between the bipolar subtypes in genetic liability for sleep traits.

## Additional Contributions

We are grateful to all participants in the Bipolar Disorder Research Network and all control participants who gave their time to be involved in this research. We are also grateful to Dr Antonio Padiñas for helpful comments on the manuscript.

## Author Contributions

ADF had full access to all the data in the study and takes responsibility for the integrity of the data and the accuracy of the data analysis.

*Study concept and design:* KJSL, ADF, LJ, NC, IJ, MOD

*Acquisition, analysis, or interpretation of data:* All authors.

*Drafting of the manuscript:* KJSL, ADF.

*Critical revision of the manuscript for important intellectual content:* KJSL, ADF, LF, KGS, HJ, IJ, LJ, SJ, NC, MOD, VEP, MNW.

*Statistical analysis:* KJSL, ADF, AR, GL, MOD

*Obtained funding:* LJ, NC, IJ, ADF

*Administrative, technical, or material support:* LF, KGS, LJ.

*Study supervision:* LJ, NC, IJ, MOD, ADF

