## Supplementary Tables and Figures for "Differences in Genetic Liability for Insomnia and Hypersomnia in Bipolar Disorder Subtypes"

### Supplemental Material

#### Supplementary Note 1

Exclusion of variants showing frequency differences between platforms PCA showed a strong effect of platform in the post-imputation dataset. To remove this effect, we performed pairwise association analyses between the three case batches, and another between the two control batches (Supplementary Table 1). Variants with association $p<0.01$ in any analysis were excluded (n=1036851). PCA performed after this step showed no effect of platform (Supplementary Figure 1).

**Supplementary Table 1.** Description of genotyping platforms for the BD and control samples. SNP N is before QC and imputation. Wave 1 of the BDRN data is part of the International Cohort Collection for Bipolar Disorder (ICCBD).

| | N cases | N controls | Chip | SNP N |
|---|---|---|---|---|
| **Wellcome Trust Case-Control Consortium (WTCCC)** | 1868 | 2934 | GeneChip 500K Mapping Array Set (Affymetrix) | 377742 |
| **Bipolar Disorder Research Network (BDRN) Wave 1** | 2577 | 2784 | Omni Express (Illumina) | 393635 |
| **Bipolar Disorder Research Network (BDRN) Wave 2** | 1104 | 0 | PsychChip (Illumina) | 578318 |

**Supplementary Figure 1.** Principal component analysis of imputed genotype data, after exclusion of variants showing platform frequency differences. Green — Wellcome Trust Case-Control Consortium samples, red— Bipolar Disorder Research Network wave 1, blue— Bipolar Disorder Research Network wave 2. See Supplementary Table 1 for descriptions of cases and controls in each sample.

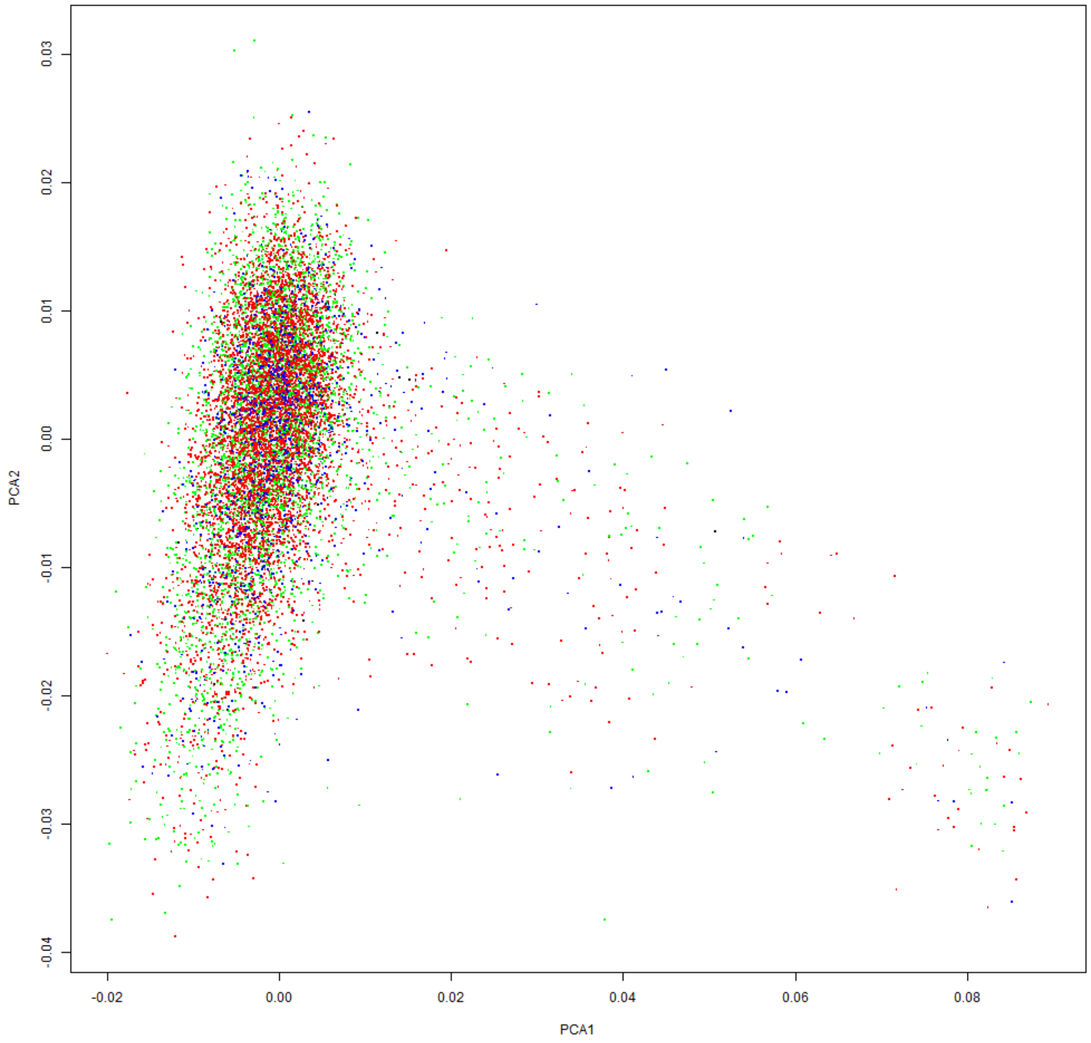

**Supplementary Table 2.** Multinomial regressions between polygenic risk scores for insomnia and bipolar subtypes compared to controls.

| | | Bipolar I disorder | | | | Bipolar II disorder | | | |
|---|---|---|---|---|---|---|---|---|---|
| PRS pT | No. SNPs | RR | 95% CI | *P*-value | *P*-value (FDR-adjusted) | RR | 95% CI | *P*-value | *P*-value (FDR-adjusted) |
| p ≤ 1 | 91950 | 0.95 | 0.91-0.99 | **0.029** | **0.044** | 1.07 | 1.01-1.14 | **0.027** | **0.044** |
| p ≤ .5 | 65942 | 0.95 | 0.91-0.99 | **0.025** | **0.044** | 1.08 | 1.01-1.15 | **0.019** | **0.044** |
| p ≤ .2 | 36927 | 0.96 | 0.92-1.01 | 0.112 | 0.130 | 1.09 | 1.03-1.17 | **0.006** | **0.028** |
| p ≤ .1 | 23718 | 0.96 | 0.92-1.00 | 0.065 | 0.091 | 1.07 | 1.01-1.15 | **0.027** | **0.044** |
| p ≤ .05 | 14652 | 0.97 | 0.93-1.01 | 0.142 | 0.153 | 1.08 | 1.01-1.15 | **0.022** | **0.044** |
| p ≤ .01 | 5415 | 0.96 | 0.92-1.01 | 0.106 | 0.130 | 1.11 | 1.04-1.18 | **0.001** | **0.009** |
| p ≤ .001 | 1410 | 0.98 | 0.94-1.03 | 0.409 | 0.409 | 1.14 | 1.07-1.21 | **8.26E-05** | **0.001** |

Analyses controlling for sex and 10 principal components. PRS pT = *P*-value threshold applied to discovery genome-wide association study in order to construct polygenic risk scores, RR = relative risk, 95% CI = 95% Confidence Interval, FDR = False Discovery Rate.

**Supplementary Table 3.** Multinomial regressions between polygenic risk scores for sleep duration and bipolar subtypes compared to controls.

| | | Bipolar I disorder | | | | Bipolar II disorder | | | |
|---|---|---|---|---|---|---|---|---|---|
| PRS pT | No. SNPs | RR | 95% CI | *P*-value | *P*-value (FDR-adjusted) | RR | 95% CI | *P*-value | *P*-value (FDR-adjusted) |
| p ≤ 1 | 92096 | 1.10 | 1.06-1.15 | **1.13E-05** | **1.07E-04** | 0.99 | 0.93-1.06 | 0.818 | 0.954 |
| p ≤ .5 | 66188 | 1.10 | 1.05-1.15 | **1.71E-05** | **1.07E-04** | 1.00 | 0.93-1.06 | 0.886 | 0.954 |
| p ≤ .2 | 37493 | 1.10 | 1.05-1.15 | **3.05E-05** | **1.07E-04** | 0.98 | 0.92-1.05 | 0.637 | 0.837 |
| p ≤ .1 | 24321 | 1.10 | 1.05-1.15 | **4.27E-05** | **1.20E-04** | 0.97 | 0.91-1.03 | 0.316 | 0.510 |
| p ≤ .05 | 15240 | 1.09 | 1.04-1.14 | **2.26E-04** | **4.52E-04** | 1.00 | 0.94-1.07 | 0.999 | 0.999 |
| p ≤ .01 | 5867 | 1.10 | 1.05-1.15 | **2.32E-05** | **1.07E-04** | 0.99 | 0.92-1.05 | 0.658 | 0.837 |
| p ≤ .001 | 1676 | 1.10 | 1.05-1.15 | **6.46E-05** | **1.51E-04** | 0.97 | 0.91-1.03 | 0.328 | 0.510 |

Analyses controlling for sex and 10 principal components. PRS pT = *P*-value threshold applied to discovery genome-wide association study in order to construct polygenic risk scores, RR = relative risk, 95% CI = 95% Confidence Interval, FDR = False Discovery Rate.

**Supplementary Table 4.** Multinomial regressions between polygenic risk scores for daytime sleepiness and bipolar subtypes compared to controls.

| | Bipolar I disorder | | | | Bipolar II disorder | | | |
|---|---|---|---|---|---|---|---|---|
| PRS pT | RR | 95% CI | *P*-value | *P*-value (FDR-adjusted) | RR | 95% CI | *P*-value | *P*-value (FDR-adjusted) |
| p ≤ 1 | 1.08 | 1.04-1.13 | **2.86E-04** | **4.55E-04** | 1.14 | 1.07-1.22 | **4.13E-05** | **1.65E-04** |
| p ≤ .5 | 1.09 | 1.04-1.14 | **2.14E-04** | **4.55E-04** | 1.14 | 1.07-1.21 | **5.79E-05** | **1.65E-04** |
| p ≤ .2 | 1.07 | 1.02-1.11 | **0.004** | **0.005** | 1.12 | 1.06-1.20 | **2.93E-04** | **4.55E-04** |
| p ≤ .1 | 1.06 | 1.01-1.10 | **0.016** | **0.017** | 1.14 | 1.07-1.22 | **4.59E-05** | **1.65E-04** |
| p ≤ .05 | 1.07 | 1.02-1.11 | **0.005** | **0.005** | 1.14 | 1.07-1.22 | **3.22E-05** | **1.65E-04** |
| p ≤ .01 | 1.09 | 1.05-1.14 | **5.89E-05** | **1.65E-04** | 1.13 | 1.06-1.20 | **2.70E-04** | **4.55E-04** |
| p ≤ .001 | 1.03 | 0.99-1.08 | 0.199 | 0.199 | 1.12 | 1.05-1.19 | **5.39E-04** | **7.55E-04** |

Analyses controlling for sex and 10 principal components. PRS pT = *P*-value threshold applied to discovery genome-wide association study in order to construct polygenic risk scores, RR = relative risk, 95% CI = 95% Confidence Interval, FDR = False Discovery Rate.

**Supplementary Table 5.** Logistic regressions between polygenic risk scores for insomnia and odds of bipolar II disorder compared to bipolar I disorder.

| PRS pT | OR | 95% CI | *P*-value | *P*-value (FDR-adjusted) | Nagelkerke $R^2$ |
|---|---|---|---|---|---|
| p ≤ 1 | 1.12 | 1.05-1.20 | **0.001** | **0.001** | 3.19E-03 |
| p ≤ .5 | 1.12 | 1.05-1.20 | **5.65E-04** | **0.001** | 3.51E-03 |
| p ≤ .2 | 1.12 | 1.05-1.20 | **6.19E-04** | **0.001** | 3.46E-03 |
| p ≤ .1 | 1.11 | 1.04-1.19 | **0.002** | **0.002** | 2.93E-03 |
| p ≤ .05 | 1.11 | 1.04-1.18 | **0.003** | **0.003** | 2.61E-03 |
| p ≤ .01 | 1.14 | 1.07-1.22 | **6.98E-05** | **2.44E-04** | 4.67E-03 |
| p ≤ .001 | 1.14 | 1.07-1.22 | **6.81E-05** | **2.44E-04** | 4.68E-03 |

Analyses controlling for sex and 10 principal components. PRS pT = *P*-value threshold applied to discovery genome-wide association study in order to construct polygenic risk scores, OR = odds ratio, 95% CI = 95% Confidence Interval, FDR = False Discovery Rate.

**Supplementary Table 6.** Logistic regressions between polygenic risk scores for sleep duration and odds of bipolar I disorder compared to bipolar II disorder.

| PRS pT | OR | 95% CI | *P*-value | P-value (FDR-adjusted) | Nagelkerke $R^2$ |
|---|---|---|---|---|---|
| p ≤ 1 | 1.11 | 1.04-1.19 | **0.002** | **0.003** | 2.71E-03 |
| p ≤ .5 | 1.11 | 1.03-1.18 | **0.004** | **0.004** | 2.50E-03 |
| p ≤ .2 | 1.11 | 1.04-1.19 | **0.002** | **0.003** | 2.91E-03 |
| p ≤ .1 | 1.13 | 1.06-1.21 | **3.20E-04** | **0.001** | 3.83E-03 |
| p ≤ .05 | 1.09 | 1.02-1.16 | **0.016** | **0.016** | 1.71E-03 |
| p ≤ .01 | 1.12 | 1.04-1.20 | **0.001** | **0.003** | 3.01E-03 |
| p ≤ .001 | 1.13 | 1.06-1.21 | **3.51E-04** | **0.001** | 3.77E-03 |

Analyses controlling for sex and 10 principal components. PRS pT = *P*-value threshold applied to discovery genome-wide association study in order to construct polygenic risk scores, OR = odds ratio, 95% CI = 95% Confidence Interval, FDR = False Discovery Rate.

**Supplementary Table 7.** Logistic regressions between polygenic risk scores for daytime sleepiness and odds of bipolar II disorder compared to bipolar I disorder.

| PRS pT | OR | 95% CI | *P*-value | *P*-value (FDR-adjusted) | Nagelkerke $R^2$ |
|---|---|---|---|---|---|
| p ≤ 1 | 1.05 | 0.99-1.12 | 0.130 | 0.182 | 6.75E-04 |
| p ≤ .5 | 1.05 | 0.98-1.12 | 0.161 | 0.188 | 5.78E-04 |
| p ≤ .2 | 1.05 | 0.99-1.13 | 0.114 | 0.182 | 7.38E-04 |
| p ≤ .1 | 1.08 | 1.01-1.15 | 0.023 | 0.082 | 1.52E-03 |
| p ≤ .05 | 1.07 | 1.00-1.15 | 0.038 | 0.090 | 1.26E-03 |
| p ≤ .01 | 1.02 | 0.96-1.09 | 0.486 | 0.486 | 1.43E-04 |
| p ≤ .001 | 1.08 | 1.01-1.15 | 0.020 | 0.082 | 1.59E-03 |

Analyses controlling for sex and 10 principal components. PRS pT = *P*-value threshold applied to discovery genome-wide association study in order to construct polygenic risk scores, OR = odds ratio, 95% CI = 95% Confidence Interval, FDR = False Discovery Rate.
